# The aging human brain exhibits reduced cerebrospinal fluid flow during sleep due to both neural and vascular factors

**DOI:** 10.1101/2025.02.22.639649

**Authors:** Sydney M. Bailes, Stephanie D. Williams, Baarbod Ashenagar, Joseph Licata, Massinissa Y. Bosli, Brandon J. Dormes, Hannah J. Yun, Dabriel Zimmerman, Alejandra Hernandez Moyers, David H. Salat, Laura D. Lewis

## Abstract

Aging reduces the quality and quantity of sleep, and greater sleep loss over the lifespan is predictive of neurodegeneration and cognitive decline. One mechanism by which sleep loss could contribute to impaired brain health is through disruption of cerebrospinal fluid (CSF) circulation. CSF is the primary waste transport system of the brain, and in young adults, CSF waves are largest during NREM sleep. However, whether sleep-dependent brain fluid physiology changes in aging is not known, due to the technical challenges of performing neuroimaging studies during sleep. We collected simultaneous fast fMRI and EEG data to measure large-scale CSF flow in healthy young and older adults and tested whether there were age-related changes to CSF dynamics during nighttime sleep. We found that sleep-dependent CSF flow was reduced in older adults, and this reduction was linked to impaired frontal EEG delta power and global hemodynamic oscillations during sleep. To identify mechanisms underlying reduced CSF flow, we used sensory and vasoactive stimuli to drive CSF flow in daytime task experiments, and found that both neural and cerebrovascular physiological changes contributed to the disruption of CSF flow during sleep. Finally, we found that this reduction in CSF flow was associated with gray matter atrophy in aging. Together, these results demonstrate that the aging human brain has reduced CSF flow during sleep, and identifies underlying neurovascular mechanisms that contribute to this age-related decline, suggesting targets for future interventions.

## Introduction

Aging significantly impacts brain health and is the primary risk factor for the development of neurodegenerative disorders. A potential contributing factor could be a change in cerebrospinal fluid (CSF) flow in the brain, since clearance of waste products and metabolites from brain tissue depends on the circulation of CSF (Iliff et al., 2012; Kress et al., 2014; Nedergaard, 2013; Tarasoff-Conway et al., 2015). A breakdown of fluid transport in the brain has therefore been proposed as a key mechanism contributing to cognitive decline and neurodegeneration in aging (Jiang-Xie et al., 2025; Lopes et al., 2024; Nedergaard & Goldman, 2020). This hypothesis has drawn support from preclinical studies showing impairment in waste clearance and drainage in animal models of aging and Alzheimer’s Disease (Kress et al., 2014; Peng et al., 2016). However, prior studies measuring CSF flow in older humans have reported conflicting results, with several observing small reductions in flow in aging (Attier-Zmudka et al., 2019; Stoquart-ElSankari et al., 2007), while others report slightly increased flow with age (Gideon et al., 1994; Vikner et al., 2024), suggesting minimal or inconsistent changes in CSF flow detected in typical daytime MRI scans in humans. It is therefore not yet clear whether typical human aging - prior to the onset of any neurological disorder - is linked to disrupted CSF dynamics.

Importantly, these prior human studies were performed during the awake state, and whether aging changes CSF flow during sleep is not known. Sleep is a state of heightened waste removal from the brain and plays a key role in maintaining brain health (Eide et al., 2021; Shokri-Kojori Ehsan et al., 2018; Xie et al., 2013). In particular, sleep reorganizes CSF circulation in the brain, driving large, low-frequency pulsatile waves of CSF flow that can be detected at the macroscopic scale (Fultz et al., 2019). In humans, CSF flow during non-rapid eye movement (NREM) sleep has been shown to be linked to coherent neural activity in the delta (0.5–4 Hz) band, also known as slow wave activity, which drives oscillations in blood volume, and in turn, drives CSF into and out of the brain through the fourth ventricle (Fultz et al., 2019). Aging is associated with significant disruptions to macroscopic sleep architecture (Carroll & Prather, 2021; Foley et al., 1995; Mander et al., 2017), along with reduced EEG slow wave activity even in non-pathological aging (Campos-Beltrán & Marshall, 2021; Landolt et al., 1996; Latreille et al., 2019; Mander et al., 2013; Ohayon et al., 2004; Varga et al., 2016). This sleep fragmentation and loss of slow waves could in theory lead to loss of CSF flow during sleep; however, whether this is true and to what extent has yet to be tested in humans.

In addition to neural slow waves, a second major factor underlying waves of CSF flow during sleep is the dynamics of the cerebrovasculature. Evidence from both rodent and human studies show that the circulation of CSF through the brain is strongly driven by vascular pulsations (Iliff et al., 2012, 2013; Kelley, 2021; Mestre et al., 2018; van Veluw et al., 2020; Vikner et al., 2024; Wang et al., 2022; Wright et al., 2024). Healthy aging significantly affects the physiology of brain blood vessels, with consequences such as stiffening (Atkinson, 2008; Cai et al., 2023; McNulty et al., 2007), and larger resting diameter that could affect the mechanical vasomotor forces that propel CSF through the brain (Juttukonda et al., 2021; Li et al., 2018). The altered vascular physiology of older individuals therefore suggests a second pathway that could influence CSF dynamics in aging.

We investigated whether and how sleep-dependent CSF flow is altered in the aging human brain. To enable simultaneous measurement of potentially contributing neural and vascular factors, we collected simultaneous EEG-fMRI-CSF measurements in healthy young and older adults during nighttime sleep. We found that older adults showed substantially reduced CSF flow during NREM sleep. This loss of CSF flow was mediated by a reduction in both frontal EEG delta power and the large hemodynamic waves that appear in youthful NREM sleep. To further probe the mechanisms that contributed to decreased CSF flow, we conducted a second set of experiments during wakefulness to causally manipulate these two potential contributors to reduced flow in aging, by driving neural activity via sensory stimulation, or vascular changes via a breath-hold task. We found significantly smaller hemodynamic responses in older adults despite similar levels of evoked neural activity, which were coupled to significantly smaller CSF flow in older adults. Together, these results show that CSF flow during sleep is reduced in the aging brain, and this decline is related to both the loss of neural frontal EEG delta power during sleep and alterations to neurovascular coupling and vascular reactivity.

## Results

To investigate how aging affects CSF flow during sleep, we collected a simultaneous EEG-fMRI-CSF dataset in a group of young (18-39 years, n=25) and older (60-84 years, n=39) subjects during nighttime rest (start time around midnight) (Fig 1A-C). This imaging method enabled simultaneous measurement of brainwide hemodynamics via the blood-oxygenation-level-dependent (BOLD) signal, and CSF flow through the fourth ventricle, representing an integrated measure of coherent fluid flow in the brain. Subjects were instructed to rest with eyes closed inside the scanner, and spontaneously transitioned into sleep during the scan, as confirmed with eye tracking videos and EEG-based sleep-staging (Berry et al., 2017).

**Figure 1.**
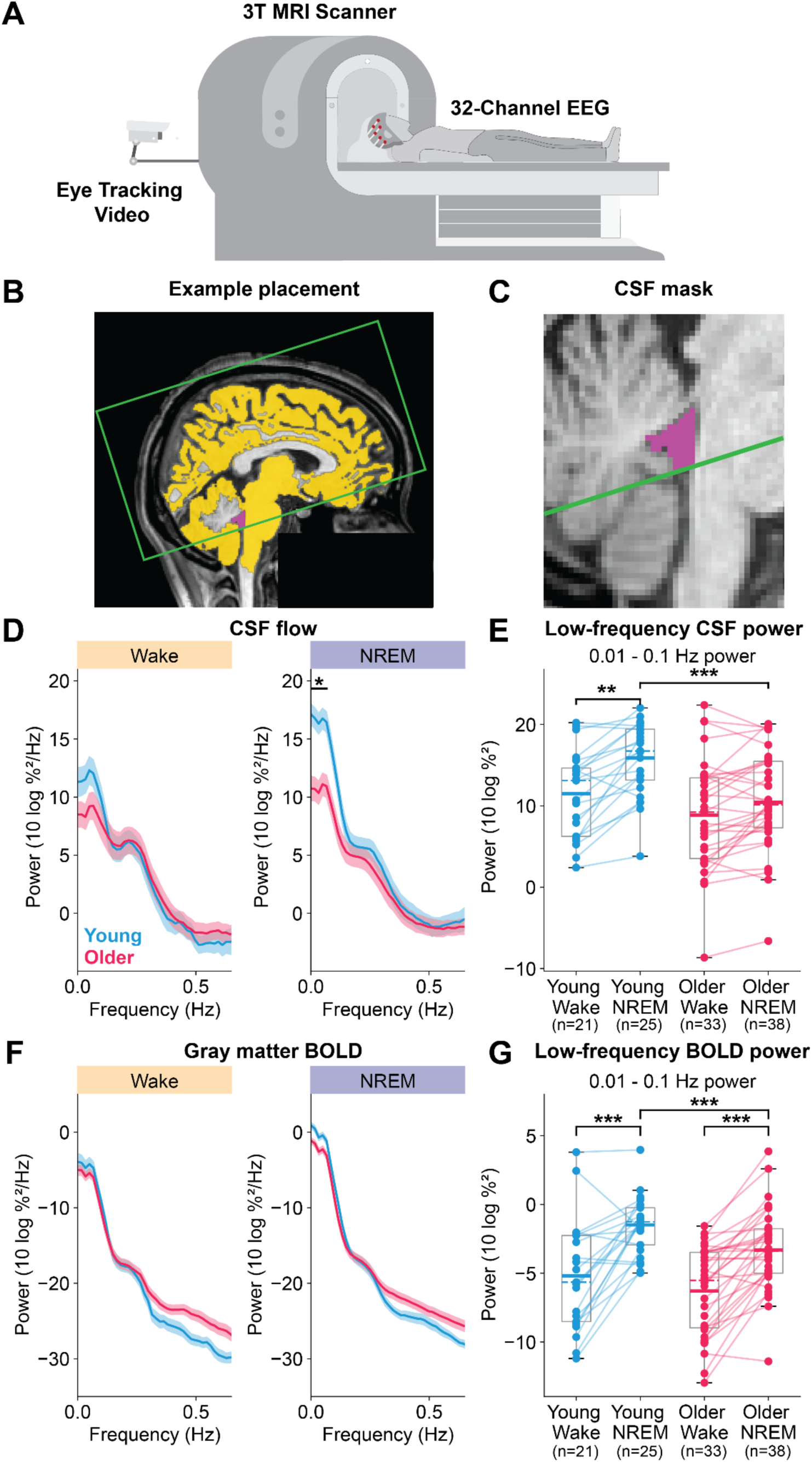
Older adults have reduced low-frequency CSF and hemodynamic oscillations during NREM sleep. **A)** We collected simultaneous EEG-fMRI data during nighttime rest along with eye-tracking video to measure whether the subject’s eyes were closed. **B)** Example positioning of fMRI acquisition volume to acquire simultaneous BOLD and CSF inflow measurements. Global gray matter is highlighted in yellow, CSF ROI is in purple, and the acquisition volume is outlined in green. **C)** Zoomed-in view of fourth ventricle and CSF mask with a green line denoting the bottom slice of the acquisition volume. **D)** CSF power spectra during wake versus during sleep show an increase in low-frequency power during NREM sleep in both groups, but a larger increase in the young adults. Young adults had significantly higher low-frequency power during NREM sleep. **E)** Total low-frequency, 0.01-0.1 Hz, power in the CSF signal significantly increases during NREM sleep compared to wakefulness in both the young and older group. Older adults have significantly reduced low-frequency CSF power during NREM sleep compared to young adults. Dashed horizontal lines represent the group median while solid horizontal lines represent the group mean. **F)** Similar to the CSF signal, low frequency gray matter BOLD power increases during NREM sleep in both groups, although the increase is smaller compared to the CSF signal. **G)** Total 0.01–0.1 Hz power in gray matter BOLD signal significantly increases during NREM sleep within both groups. During NREM sleep, older adults have significantly reduced low-frequency hemodynamic oscillations.

Because large low-frequency (∼0.05 Hz) waves of CSF flow appear during NREM sleep in young adults (Fultz et al., 2019) we first examined the frequency content of the CSF flow signal across sustained periods of wakefulness and NREM sleep (minimum duration=60 seconds). We found that during wakefulness there were no group differences in the CSF flow power spectrum, but during NREM sleep the young adults had significantly higher low-frequency CSF power compared to older adults (Fig 1D, p<0.05 for [0, 0.066] Hz, Wilcoxon rank-sum test, Holm-Bonferroni corrected). Low-frequency CSF power increased during NREM sleep compared to wakefulness in young adults, consistent with prior work, and analyzing CSF flow bandpower specific to the 0.01-0.1 Hz range confirmed that young adults show a significant increase in low-frequency (0.01-0.1 Hz) power during NREM sleep (Fig 1E, p=0.008, Wilcoxon rank-sum test). However, we did not find a significant effect of NREM sleep on low-frequency CSF power in older adults (Fig 1E, p=0.254, Wilcoxon rank-sum test). While the group-level difference in the older adults between wakefulness and NREM sleep was not significant, not all subjects had sustained periods of both wakefulness and NREM sleep within a session. To assess if a more subtle change in CSF flow was present on an individual level, we performed a within-subject analysis, including only subjects with both wakefulness and sleep within a scan session. This paired analysis showed that both young individuals and older individuals showed increased low-frequency CSF power during NREM sleep (Supp. Fig. 1A), but the magnitude of the sleep-related increase in low-frequency CSF power was significantly lower in older adults (young average change=4.00 dB, older average change=1.65 dB, p=0.02, Wilcoxon rank-sum test). Low-frequency 0.1-1 Hz CSF power was also significantly lower in older adults during NREM sleep, as compared to young adults (p<0.001, Wilcoxon rank-sum test). We next constructed a linear model to examine the interaction of age and sleep (CSF power∼Age*State*Sex) and found that aging specifically reduced the sleep-dependent increase in CSF flow (p_Age:State_=0.005, linear mixed effects model). Because aging is known to alter the total volume of CSF, we also confirmed that these differences in CSF flow signals were not due to confounding differences in the cross-sectional area or volume of the fourth ventricle, where flow was measured (Supp. Fig 2). Taken together, these results demonstrated that low-frequency, sleep-dependent CSF flow was significantly reduced in aging.

A key driver of CSF flow waves is coupled waves in global hemodynamics, because changes in cerebral blood volume draw CSF into and out of the skull (Mokri, 2001). In young adults, CSF oscillations are strongly anticorrelated with hemodynamic oscillations measured using BOLD fMRI (Fultz et al., 2019), so we next examined if the hemodynamic signal showed similar state- and age-dependent changes as the CSF flow signal. We found that, similar to CSF, low-frequency global gray matter BOLD power significantly increased from wakefulness to sleep in young adults (Fig 1G, p<0.001, Wilcoxon rank-sum test). Older adults also exhibited a significant increase in BOLD fluctuations from wakefulness to sleep (Fig 1G, p<0.001, Wilcoxon rank-sum test), but their low-frequency gray matter BOLD power during NREM sleep was significantly lower than young adults (Fig 1G, p_sleep_=0.002, Wilcoxon rank-sum test). Similar to the CSF flow results, the effect of aging on hemodynamics was significantly stronger during sleep, compared to wakefulness (mixed-effects model: Gray matter BOLD power∼Age*State*Sex, p_Age:Stage_=0.013).

These results demonstrated that older adults have dampened effects of sleep on both low-frequency CSF flow and hemodynamic waves. We next examined if and to what extent the coupling between CSF and hemodynamic oscillations are impaired in older adults. We found that the sleep-related increase in low-frequency CSF power was significantly correlated with the increase in low-frequency BOLD power within both groups (Fig. 2A), suggesting that coupling between large-scale hemodynamic and CSF oscillations is mostly maintained with age. CSF typically shows strong cross-correlation with the negative derivative of the gray matter BOLD signal, consistent with an alternation of changes in blood volume and CSF flow (Fultz et al., 2019). To assess this coupling, we calculated this cross-correlation during NREM sleep in both age groups. Consistent with prior studies, in young adults, CSF flow waves were significantly and strongly anti-correlated with gray matter BOLD oscillations (Fig 2B, maximal r=0.79, at lag -1.13 s, p<0.001, permutation test). The older adults similarly exhibited significant correlation between BOLD and CSF oscillations (Fig 2B, maximal r = 0.68 at lag -0.76 s, p<0.001, permutation test), with a small but significant reduction in the magnitude of the correlation (difference=0.11; p=0.008, permutation test). We further confirmed that differences in the strength of coupling were not significantly related to motion confounds (Supp. Fig 4A). These results demonstrated that reduced CSF flow was tightly associated with reduced hemodynamic waves during sleep in both groups, with only slightly weakened hemodynamic-CSF coupling in the older group.

**Figure 2.**
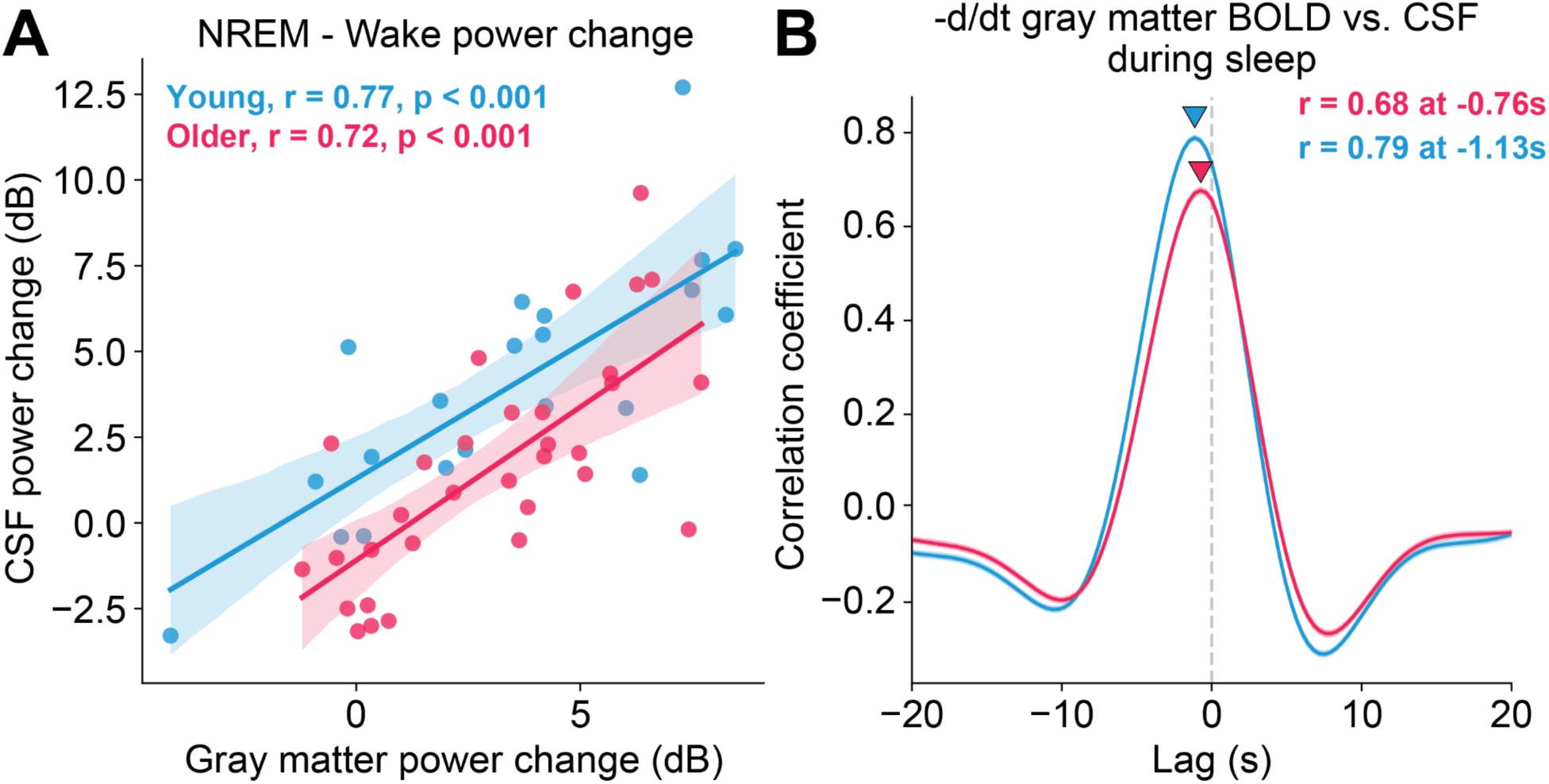
Coupling between hemodynamic and CSF oscillations is significantly weaker in older adults. **A)** In both young and older adults, there is a significant correlation between the change in low-frequency gray matter BOLD power and CSF power. **B)** In both groups, CSF is strongly correlated with the zero-thresholded negative derivative of gray matter BOLD, with slightly lower correlation in older adults.

Multiple potential mechanisms could lead to reduced hemodynamic and CSF waves during NREM sleep. In young adults during NREM sleep, CSF waves are coupled to EEG delta (0.05–4 Hz) power (Fultz et al., 2019), and coherent neural activity can induce CSF flow (Holstein-Rønsbo et al., 2023; Williams et al., 2023). Therefore, we hypothesized that a reduction in EEG delta power during NREM sleep in older adults could contribute to reduced CSF flow. To examine neural differences across groups, we extracted EEG delta power (Fig. 3A), focusing on the frontal regions due to their well-known concentration of delta power during sleep (Finelli et al., 2001; Werth et al., 1997). Frontal EEG delta power was calculated from channel Fz (except in subjects or runs with bad-quality data on that channel, in which case we used F3, n_subjects_=7, n_runs_=10). On an individual level, both groups exhibited an increase in frontal EEG delta power from wakefulness to sleep (p_young_<0.001, p_older_<0.001, Wilcoxon rank-sum test). Consistent with prior work (Landolt et al., 1996; Latreille et al., 2019; Varga et al., 2016), we found that older adults had significantly reduced frontal EEG delta power compared to young adults, both during wakefulness and, more prominently, during NREM sleep (Fig 3B, p_sleep_<0.001, p_wake_=0.018, Wilcoxon rank-sum test). Similar to both the CSF flow and gray matter BOLD results, the effect of aging on frontal EEG delta power was significantly stronger during sleep compared to wakefulness (mixed-effects model: frontal EEG delta power∼Age*State*Sex, p_Age:Stage_=0.049). To test if the loss of sleep-specific frontal EEG delta power was related to CSF flow, we calculated the correlation between the change in frontal EEG delta power and the change in low-frequency CSF power within the two groups. Frontal EEG delta power was strongly correlated with CSF flow in young adults (Fig 3C, r=0.73, p<0.001, Wald test): individuals with strong frontal EEG delta power during sleep also exhibited the greatest increase in CSF flow. However, in older adults, the increase in frontal EEG delta power was trending but not significantly associated with CSF flow (Fig 3C, r=0.35, p=0.052, Wald test) and this relationship was significantly weaker in older adults than in young adults (p=0.039, Fisher z-transformation). To further investigate the coupling between frontal EEG delta and CSF we then tested whether the temporal relationship between the signals differed in the two age groups. We found that both groups had a strong relationship between the EEG and CSF signals, with EEG leading CSF (Fig 3D, maximal r_young_=0.27 at 8.32s, maximal r_older_=0.14 at 9.23s), but this coupling was significantly weaker in older adults (difference=0.13, p<0.001, permutation test). We additionally confirmed these differences were not due to motion confounds (Supp. Fig 4B). We hypothesized that the reduced correlation between EEG and CSF in older adults was at least partially due to the reduced amplitude and low temporal variability in the frontal EEG delta activity in older adults. To test this theory we examined frontal EEG delta activity locked to individual peaks in large CSF waves. We found that not all older adults had identifiable peaks in their CSF waves (n_young_=22/25, n_older_=27/39), but in individuals that did exhibit CSF peaks, frontal EEG delta activity was lower-amplitude and exhibited a less prominent peak compared to the young subjects (Fig 3E, nCSFPeaks_young_=869, nCSFPeaks_older_=457, 5.87μV difference, p<0.001, permutation test). These results demonstrated that reduced frontal EEG delta power during NREM sleep was associated with loss of CSF flow, and that older adults have impaired coupling between frontal EEG delta and CSF flow in part due to a significant loss of the neural driver of CSF flow.

**Figure 3.**
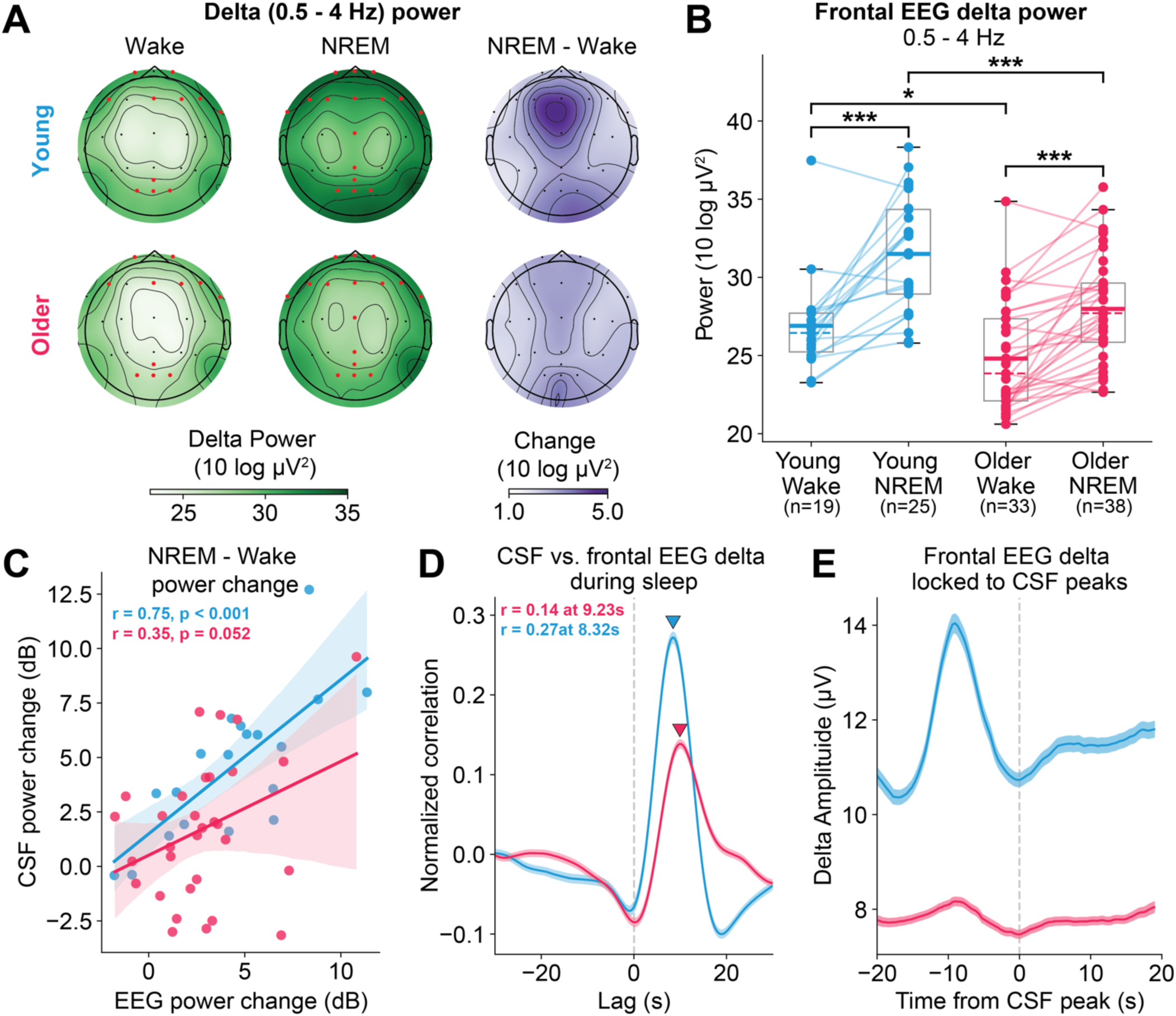
Older adults have less EEG delta power and weaker coupling between EEG and CSF. **A)** (Top row) Topographical head maps of delta (0.5–4 Hz) power in young subjects during wake (left) and NREM sleep (middle), and the increase in power from wakefulness to NREM sleep (right). (Bottom row) Topographical head maps of delta power in older subjects during wake (left) and NREM sleep (middle), and the increase in power from wakefulness to NREM sleep. Young subjects have significantly higher delta power during both wake and NREM sleep across frontal and occipital electrodes (significant electrodes marked in red). The strongest changes during NREM sleep were focused in the frontal electrodes while in older adults the changes are weaker and more distributed across the scalp. **B)** The magnitude of frontal EEG delta power significantly increases during NREM sleep compared to wakefulness in both groups. Older adults have significantly less frontal EEG delta power during NREM sleep compared to young adults, and frontal EEG delta power during sleep is significantly more impacted by age compared to wake frontal EEG delta power. **C)** In young adults, the change in frontal EEG delta power from wake to sleep is significantly and strongly correlated with the change in low-frequency CSF power; however, in older adults the relationship is non-significant and significantly weaker. **D)** The correlation between frontal EEG delta and CSF oscillations is significantly stronger in young adults compared to older adults. **E)** The frontal EEG delta envelope locked to individual waves of CSF is significantly reduced in amplitude in older adults compared to young adults.

A second factor that could contribute to reduced CSF flow is altered vascular physiology, because the vasculature is a key driver of CSF flow, both due to neurovascular coupling as well as systemic vasoconstriction (Fultz et al., 2019; Picchioni et al., 2022; van Veluw et al., 2020; Wang et al., 2022). To examine the role of neural and vascular drivers of CSF flow in aging, we designed a serial mediation model with age group as the predictor variable (X) and low-frequency CSF power during NREM sleep as the dependent variable (Y, Fig 4). Large-scale coherent neural activity in the delta power band is known to modulate blood volume which in turn drives CSF flow during sleep (Fultz et al., 2019). Therefore, the first mediator in the sequence was frontal EEG delta power during NREM sleep (M_1_) followed by gray matter BOLD low-frequency power during NREM sleep (M_2_). The analysis revealed a significant total effect of age on CSF power (c=5.46, p<0.001, [2.77, 8.17], 95% CI) but a non-significant direct effect of age on CSF power when controlling for the mediators (c’=2.20, p=0.107, [-0.476, 4.88]) suggesting that the entire effect of age group on CSF flow during sleep can be explained by these neural and hemodynamic mediating processes. Our analysis showed significant paths from age group to frontal EEG delta power (a_1_=2.85, p<0.001, [1.59, 4.11]) and from frontal EEG delta power to gray matter BOLD low-frequency power (b_2_=0.635, p=0.008, [0.167, 1.10]). The mediation path from age group to low-frequency BOLD power, however, was not significant (a_2_=0.949, p=0.182, [-0.44, 2.34]). Together these results demonstrated that frontal EEG delta power mediates the relationship between age group and CSF low-frequency power both directly and through low-frequency gray matter BOLD power, indicating that both neural and hemodynamic factors contribute to loss of CSF flow in aging.

**Figure 4.**
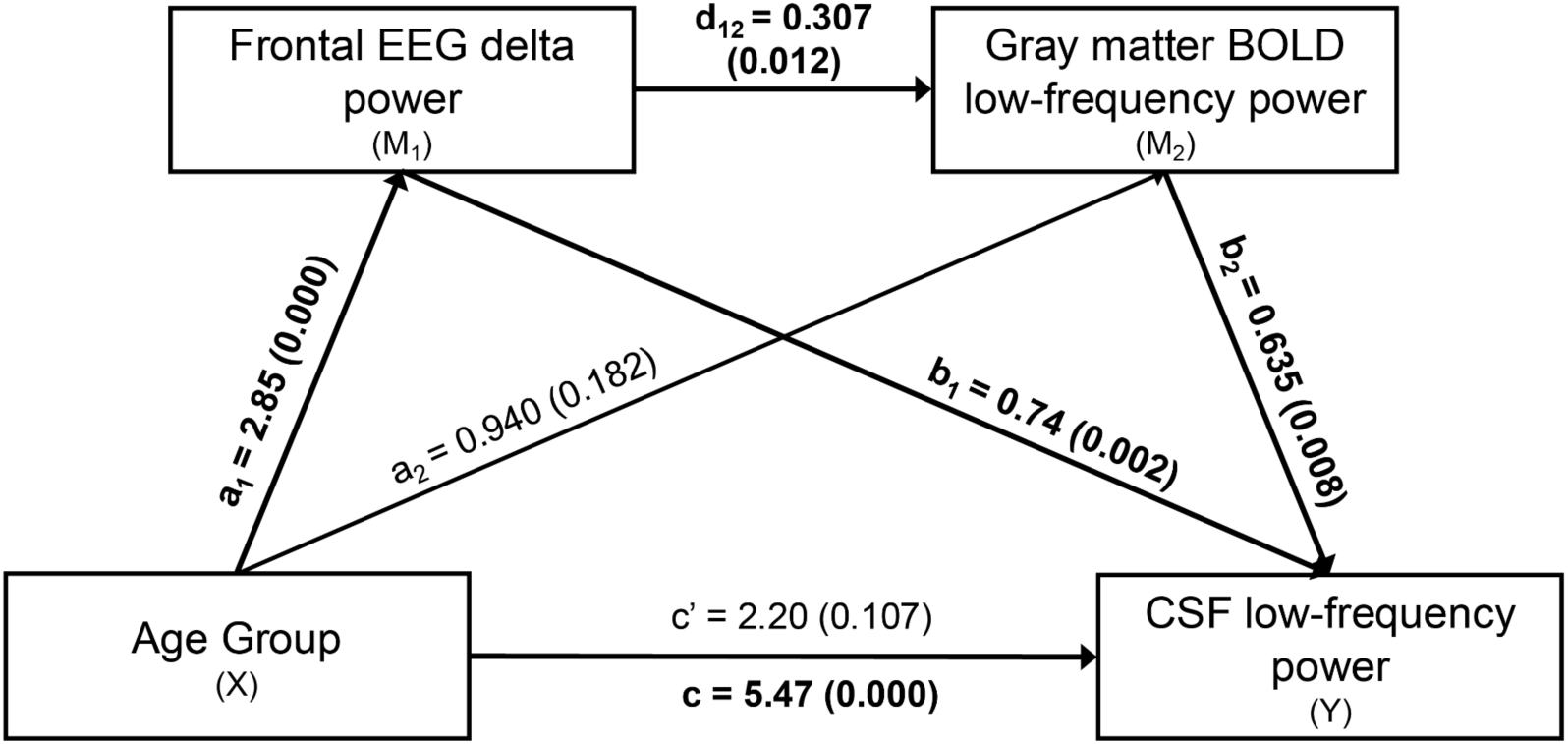
The reduction in low-frequency CSF power during sleep in the older age group is mediated by frontal EEG delta power both directly and through its effect on global gray matter BOLD low-frequency power. There is a significant total effect of age group on CSF low-frequency power; however, when accounting for the mediators, the relationship is no longer significant indicating full mediation. There were significant paths from age group to frontal EEG delta power, from frontal EEG delta power to gray matter BOLD power, and gray matter BOLD to CSF power . However, the direct path from age group to low-frequency gray matter BOLD power is not significant.

These statistical associations suggested that two mechanisms contribute to decreased CSF flow in aging: loss of neural activity in the delta power band associated with slow waves, and reduced vascular reactivity. We therefore performed a second experiment to causally manipulate neural activity and vascular physiology, hypothesizing that reduced neurovascular coupling contributed to reduced CSF flow. We conducted a second daytime study with EEG-fMRI using controlled stimuli that are known to drive CSF flow (n=31 young adults; n=55 older adults). We first tested whether there were cerebrovascular reactivity (CVR) differences between the young and older groups, as prior work has shown that CVR declines with age (Catchlove et al., 2018; Gauthier et al., 2013; Liu et al., 2013; Miller et al., 2019). To drive vascular responses, we used a 15-second breath hold task (Fig 5A), and nasal cannula to measure expired CO_2_ during the task (Fig 5B), to directly dilate blood vessels via changes in blood oxygenation, and obtained usable measurements in a subset of the cohort (n_older_=31, n_young_=19; see Methods).

**Figure 5.**
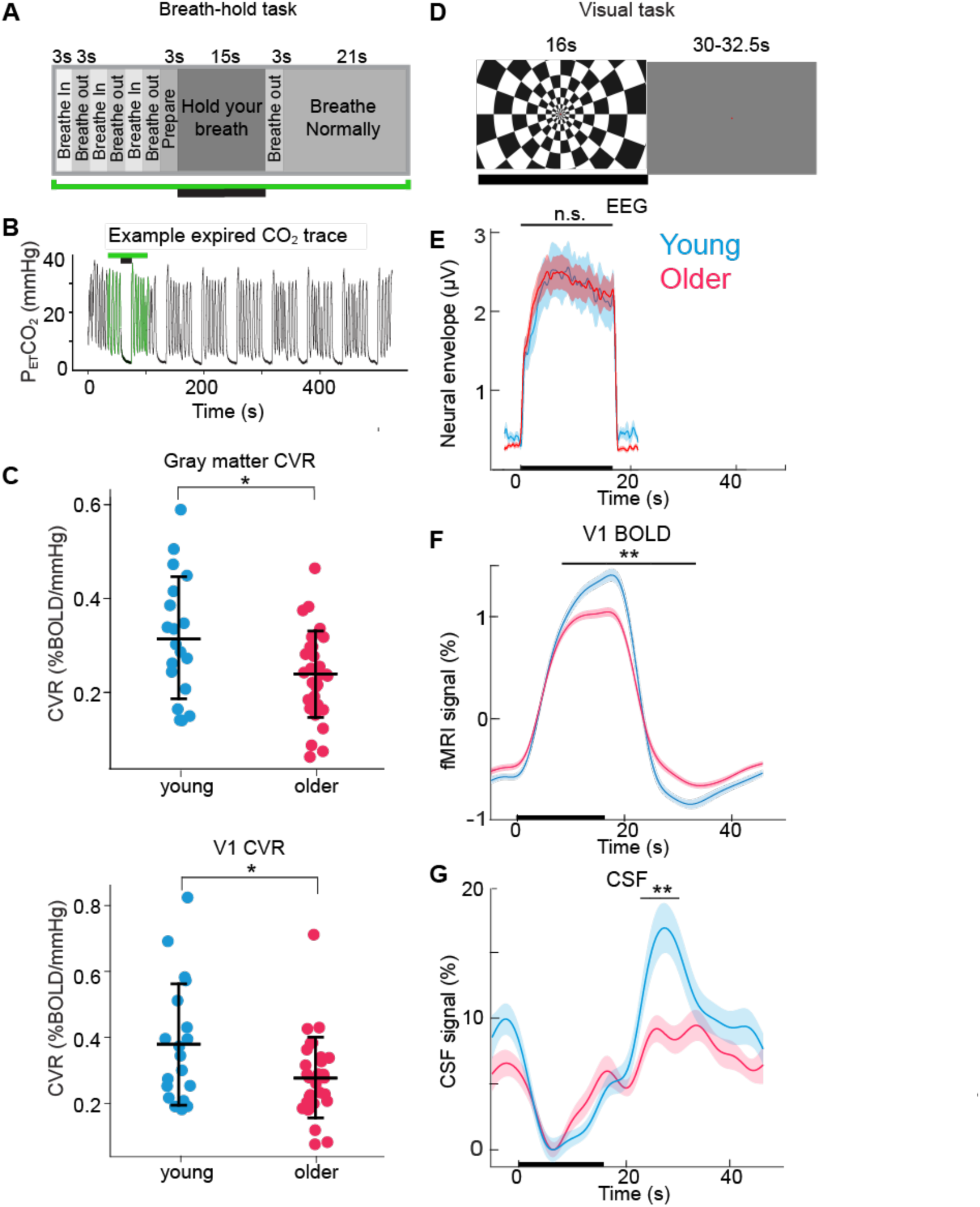
Daytime awake experiments using controlled stimuli to drive neural and vascular responses show that cerebrovascular reactivity and neurovascular coupling-driven CSF flow are reduced in older adults. **A)** Schematic of the breath-hold task used to assess cerebrovascular reactivity. Expired CO_2_ was measured via a nasal cannula and was used for quantitative cerebrovascular reactivity (CVR) mapping. **B)** Example end-tidal CO_2_ trace measured via a nasal cannula for one task run. Black bars indicate 15-second breath-hold periods. **C)** CVR was lower in older adults (red) than in young adults (blue) in all gray matter and in visual cortex. **D)** Schematic of the flickering 12 Hz visual checkerboard task designed to assess neurovascular coupling during the awake state. ON blocks consisted of an inverting 12 Hz stimulus (16s) while OFF blocks consisted of a gray screen (jittered timing, 30 - 32.5 s). **E )** Waveforms are the mean group-level evoked neural responses from the average of four occipital EEG channels are displaced. The neural responses did not differ between older and young groups. Time 0 s indicates the onset of the ‘ON’ blocks. Shading reflects standard error across subjects. **F)** The mean V1 BOLD response was significantly smaller in older than in younger subjects during t = [8.3 s to 33.2 s] (p<0.01, Wilcoxon sign rank test, Bonferroni-Holm corrected for multiple time bins). Shading is standard error. **G)** Task-evoked CSF flow was significantly reduced in older adults relative to younger adults during t =[24.9 to 33.2 s] (p<0.01, Wilcoxon sign rank test, Holm-Bonferroni corrected for multiple time bins). Stars indicate significance: * <0.05, ** <0.01.

We found that older adults had lower levels of CVR both globally across gray matter (p=0.018, two-sample t-test) and in primary visual cortex specifically (p=0.025, two-sample t-test) relative to younger adults (Fig 5C), consistent with prior studies (Catchlove et al., 2018; Li et al., 2018; Miller et al., 2019).

To test whether neurovascular coupling differed between young and older groups, we used a visual stimulus that is known to drive substantial changes in cerebral blood volume (Ciris et al., 2014; Donahue et al., 2009) and measured the evoked neural (EEG), hemodynamic (BOLD), and CSF signals in response to the task (Fig 5D). Previous work has established that sensory-evoked neural activity can drive waves of cerebrospinal fluid flow via changes in blood volume due to neurovascular coupling (Williams et al., 2023). We tested whether there were differences in this neural drive of CSF flow in older adults by inducing neural activity using sensory stimulation. Subjects viewed a checkerboard stimulus flickering at 12 Hz in a block design (16-second ON block, 30-32.5- second OFF block). Aging can cause deterioration of visual perception and contrast sensitivity (Zhuang et al., 2021), so we used strict exclusion criteria to select subjects with good vision. In addition, to control for possible differences in contrast sensitivity, we tested two stimulus contrast levels to identify a very high-contrast stimulus that saturated the maximal response in both groups (Supp. Fig 5, p>0.05 for all time bins). To measure the evoked neural response, we calculated the mean stimulus-evoked neural activity of four occipital electrodes (Pz, O1, O2, Oz) and extracted the amplitude envelope between 11.9-12.1 Hz. We observed clear evoked neural responses in both the young (n=28, 3 subjects excluded for unusable EEG data) and older groups (n=54, 1 subject excluded for unusable EEG, Fig 5E), and the amplitude of stimulus-evoked neural activity did not significantly differ between older and younger adults (Fig 5E, Wilcoxon rank-sum test, p>0.05 for all time bins).

Next, we tested whether evoked BOLD responses differed between young (n=31) and older (n=55) groups, as prior work has suggested hemodynamic responses to visual stimuli are reduced in older adults (West et al., 2019). In contrast with the equivalent stimulus-evoked neural activity across age groups, we observed significant age-related reductions in stimulus-evoked BOLD response in primary visual cortex (V1, Fig 5F, p<0.01 for t =[8.3, 33.2] s, Wilcoxon rank sum test, Holm-Bonferroni corrected). In addition to reduced responses in V1, older adults also had significantly lower global gray matter BOLD responses than young adults (p=0.036, unpaired t-test), suggesting that there was also widespread reduction in BOLD responses across the brain. The reduced V1 and global gray matter response in older adults suggested that aging reduced neurovascular coupling, even when neural activity was held constant across the young and old groups.

We next investigated the consequences of reduced hemodynamic responses in the older group on the magnitude of evoked CSF flow. We hypothesized that stimulus-evoked CSF flow would be maximal during the period when V1 and gray matter BOLD signals declined, because the blood and CSF fluid compartments should alternate due to volume displacement, leading to upwards CSF flow after stimulus offset (Williams et al., 2023). Analogous to the BOLD results, we found a significant reduction in evoked CSF flow in older adults relative to young adults (Fig 5G, t = [24.9, 33.2] s, p<0.01, Wilcoxon signed rank test, Holm-Bonferroni corrected). This result showed that reduced stimulus-evoked BOLD hemodynamics were associated with significantly reduced CSF flow, despite equivalent neural activity, suggesting that reduced vascular responses contribute to reduced CSF flow in older adults.

Finally, we examined whether CSF flow was related to anatomical brain changes in older adults, as cortical gray matter atrophy increases with age (Salat et al., 2004). We first calculated the gray matter volume in the frontal lobe, because EEG delta power reductions are typically found in frontal regions, in the literature (Finelli et al., 2001; Werth et al., 1997), as well as in our data. Frontal lobe gray matter volume (normalized to total intracranial volume) was significantly lower in the older adults (Fig 6A, p<0.001, Wilcoxon rank-sum test), as expected. We found that normalized frontal gray matter volume was significantly correlated with NREM CSF power in older adults (Fig. 6B, r_older_=0.35, p_older_=0.03). The same relationship within the young population was non-significant (Fig. 6B; r_young_=- 0.13, p_young_=0.551) possibly due to less inherent variation in both the functional and structural measures of interest in young adults. Additionally, normalized gray matter volume in other lobes was not significantly associated with NREM CSF power (Supp. Fig 6), suggesting that the maintenance of frontal structural integrity, in particular, is important for CSF flow.

**Figure 6.**
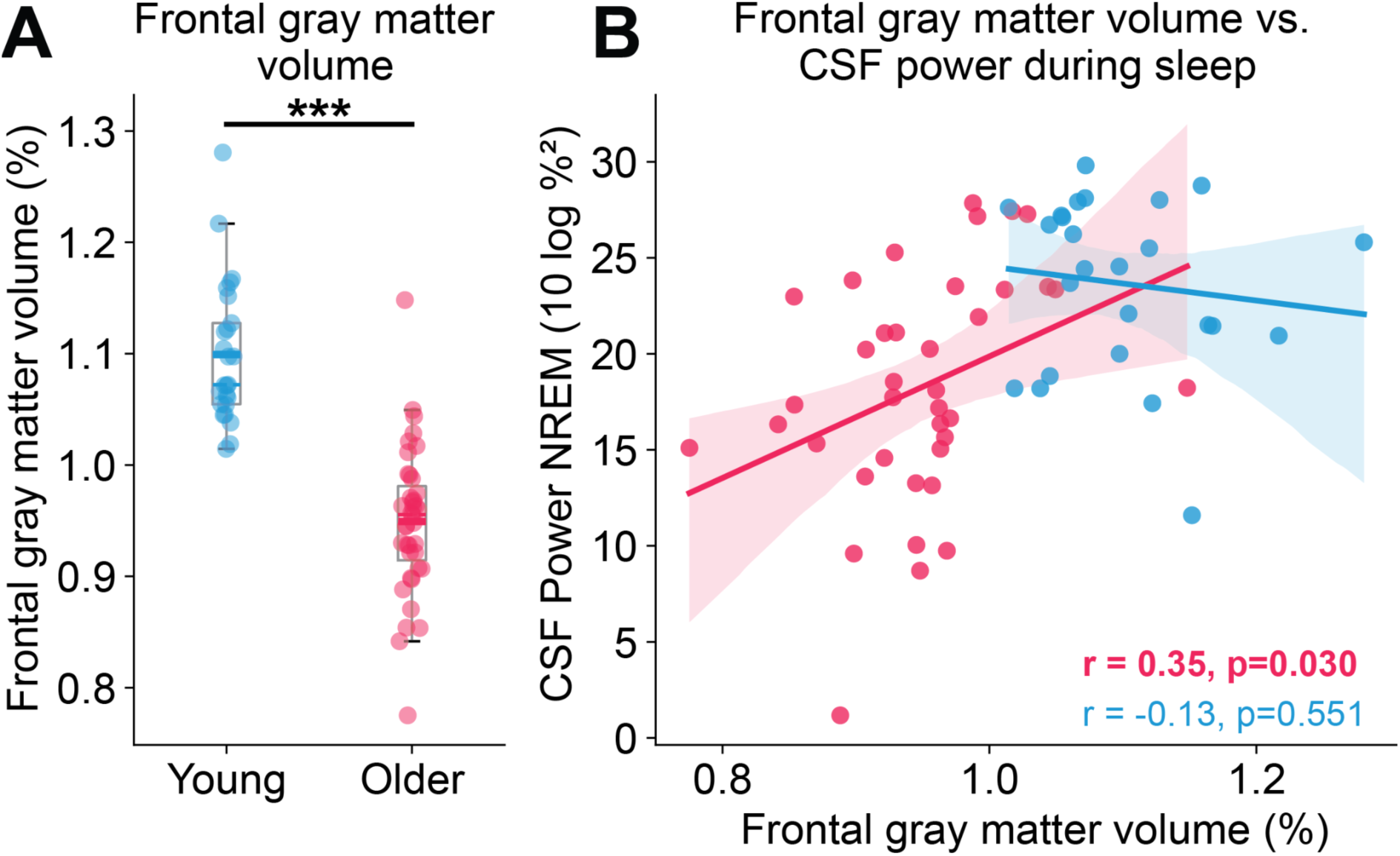
CSF low-frequency power and strength of coupling between CSF and EEG are significantly associated with frontal gray matter volume in older adults. **A)** Older adults have significantly less frontal gray matter volume normalized to total intracranial volume. **B)** Low-frequency CSF power during NREM sleep is significantly correlated with frontal lobe gray matter volume normalized to total intracranial volume only in older adults.

## Discussion

We conclude that in the aging human brain, the hemodynamic and CSF flow waves that appear during NREM sleep are substantially reduced. CSF flow is strongly driven by vascular fluctuations, which can also be driven by neural activity, and our results demonstrate that both these neural and vascular mechanisms contribute to the loss of CSF flow in aging. Older adults had reduced EEG delta power during sleep and impaired coupling between neural, hemodynamic, and CSF oscillations. Furthermore, daytime tasks causally driving CSF flow via neural activity and manipulating the cerebrovasculature revealed impaired neurovascular coupling and vascular reactivity. The reduced CSF flow we observed during sleep was further associated with structural brain changes, as smaller CSF flow magnitude was associated with greater frontal gray matter atrophy. These results show that sleep-dependent CSF flow is reduced in the aging brain, and identifies both neural and vascular mechanisms contributing to this impairment.

In agreement with prior work, we found that EEG power in the slow wave activity band during NREM sleep is significantly lower in older adults compared to young adults (Campos-Beltrán & Marshall, 2021; Landolt et al., 1996; Landolt & Borbély, 2001; Mander et al., 2013; Ohayon et al., 2004; Varga et al., 2016). Additionally, we found that this reduction in slow wave power is the strongest mediator of loss of NREM CSF flow, both directly and indirectly through its effect on global hemodynamics. The relationship between EEG and hemodynamics is affected by multiple factors including cerebrovascular reactivity and neurovascular coupling. While there is general agreement in the literature that cerebrovascular reactivity declines with age (Catchlove et al., 2018; Miller et al., 2019), neurovascular coupling has mixed results. Some studies have suggested that neurovascular coupling is not significantly impacted by normal aging (Grinband et al., 2017; Rosengarten et al., 2003) while others report impaired neurovascular coupling in aging (D’Esposito et al., 2003; Lipecz et al., 2019) . Our reported results are consistent with the idea that neurovascular coupling is reduced in healthy aging since we saw that the same level of sensory-evoked neural activity resulted in significantly reduced hemodynamic responses in older adults. Furthermore, loss of slow wave power mediates the reduction of low-frequency CSF pulsations in older adults through its effect on global hemodynamics, consistent with the idea that neurovascular coupling impairments result in reduced low-frequency hemodynamic oscillations which in turn drives less CSF flow. Ultimately, this work demonstrates that, while changes in neural activity driving CSF flow during sleep are profound, changes to the cerebrovasculature are also important factors influencing CSF flow. Future work exploring CSF flow through fluid compartments in the whole brain, including lateral ventricles and perivascular spaces, would further elucidate the relationship between the cerebrovasculature and fluid transport.

We found that the maintenance of frontal lobe structural integrity, in particular, is important for CSF flow during sleep in the aging brain. Frontal brain regions have been linked to the generation and propagation of large-scale coherent neural slow waves in the delta band that are known to drive oscillations in blood volume and ultimately CSF flow in and out of the brain through the fourth ventricle during sleep (Fultz et al., 2019; Murphy et al., 2009). One possibility is that the loss of frontal gray matter volume is associated with reduced low-frequency CSF flow due to a loss of neural slow waves that contribute to CSF flow during sleep. This is consistent with evidence linking structural brain changes in medial prefrontal cortex to reduced slow wave activity in older adults (Dubé et al., 2015; Latreille et al., 2019; Mander et al., 2013); however, more work is needed to determine the precise causal mechanisms.

Gray matter atrophy, and brain aging in general, is highly variable across individuals. While we observed large group-level differences in CSF flow and the effects of sleep, there was also substantial subject-level variability with some older subjects exhibiting more “young-like” traits. For example, some older adults showed similar levels of low-frequency CSF power during NREM sleep to the young population, and some older adults had comparable CVR to young adults. We also found that there was subject-level variability in the strength of coupling between EEG, BOLD hemodynamics, and CSF. The interactions between these systems ultimately dictate the amount of CSF flow during NREM sleep. For example, two subjects with similar levels of frontal EEG delta power during sleep but different levels of impaired neurovascular coupling would have different levels of fluid flow during sleep, as slow waves would not be able to recruit hemodynamic oscillations and driving CSF flow as strongly compared in individuals without functionally intact cerebrovasculature. This variance suggests that aging may impair CSF flow in different individuals via different mechanisms, and that individual-level imaging may be needed to identify which factor is leading to altered aging CSF circulation. Importantly, while in some individuals loss of sleep and slow waves are a critical determinant of CSF flow, other individuals may have adequate sleep and nevertheless experience impaired CSF flow due to impaired vascular reactivity. These distinct mechanistic origins would in turn suggest distinct therapeutic interventions for different individuals.

Circulation of CSF is critical for the removal of waste products and metabolites from brain tissue, and most studies have demonstrated that it is particularly heightened during sleep (Eide et al., 2021; Shokri-Kojori Ehsan et al., 2018; Xie et al., 2013), although see (Miao et al., 2024, Kroesbergen et al., 2024). Our results demonstrate that there are significant disruptions to macroscopic CSF flow in the fourth ventricle during sleep in typical aging, prior to any diagnosed neurological disorder. This ventricle-level CSF flow measurement indicates that global, integrated CSF movement is impaired in older adults. The functional consequences of reduced large-scale ventricular CSF flow for waste clearance are still not established. A reduction in the pulsatile vascular dynamics driving CSF flow in older adults would be expected to negatively impact CSF-interstitial fluid (ISF) exchange and reduce molecular clearance from brain tissue (Iliff et al., 2013; Mestre et al., 2018). However, this study could not directly measure tracer clearance, as it requires invasive procedures. Previous studies have demonstrated an association between macroscopic CSF flow and clinical outcomes: lower CSF flow is linked to worse cognitive performance across multiple domains (Attier-Zmudka et al., 2019), and patients with Alzheimer’s disease pathologies have significantly reduced macroscopic cerebrovascular-CSF and global BOLD-CSF coupling during rest (Ferdinando et al., 2023; Han et al., 2021) Still, future work should determine whether and to what extent the reduced macroscopic CSF flow identified during sleep in aging is linked to higher protein accumulation (e.g. tau and amyloid) or impaired cognitive function to clarify how CSF flow during sleep impacts clinical outcomes. Recent work using invasive intrathecal injections in human patients has demonstrated success in tracking clearance (Eide et al., 2021), suggesting an avenue for future research to directly test the relationship between large-scale CSF flow during sleep and clearance from the brain. In addition, our results demonstrated reduced global hemodynamic waves in older adults as well, suggesting a loss of the critical vascular pulsations needed to drive transport at the perivascular scale.

Mounting evidence in the literature indicates that poor sleep is predictive of cognitive decline and neurodegeneration across the lifespan (Ibrahim et al., 2024; Lucey et al., 2019; Sabia et al., 2021), but the mechanisms underlying this relationship are unknown. An impaired waste clearance system has been proposed as one pathway by which sleep impacts brain health (Jiang-Xie et al., 2025; Nedergaard & Goldman, 2020). Our work shows that CSF flow is reduced during sleep in healthy older adults before the onset of any neurological or neurodegenerative diseases; intriguingly, our method also shows high variance across older adults, with some older individuals exhibiting relatively high vs. low CSF flow. Future work should investigate the clinical utility of assessing CSF dynamics for early detection of neurodegeneration.

Together, our results show that aging leads to the deterioration of multiple drivers of CSF flow: both neural slow waves during sleep, and the vascular fluctuations that propel CSF flow. These age-related changes contribute to reduced circulation of CSF in the aged brain, particularly during sleep, but also provide targets for future interventions.

## Methods

### Subject population and exclusion criteriah

Data acquisition was performed at two sites. All subjects provided written informed consent. At Site 1, procedures were approved by the Boston University Charles River Campus Institutional Review Board (IRB #6031). At Site 2, procedures were approved by the Massachusetts Institute of Technology Institutional Review Board (IRB #23030000951). Exclusion criteria for the whole study included the regular use of sleep aid medications, current psychiatric or neurological disorders, cognitive impairment, major medical disorders, or sleep disorders, substance use disorders or excessive (estimated >=400 mg daily) caffeine intake, or any MRI contraindications. Both a nighttime sleep experiment and daytime experiment were performed at both sites. Subjects recruited for the older adult group were between ages 60 and 85. Subjects recruited for the young adult group were between ages 18 and 40.

For the nighttime sleep experiment, 70 subjects in total completed an MRI session (Site 1: n=20 young, n=22 older; Site 2: n=8 young, n=20 older). Night scans were scheduled to begin around midnight, with the exact start time depending on factors related to the equipment set up and preparation on EEG caps. Subjects were instructed not to consume any caffeine after 11 am on the day of the experiment and not to nap the day of the experiment. Each subject was scanned for 1-3 runs of sleep opportunity, with each run lasting 25 minutes. The number of runs depended on factors such as experimental setup time and subject comfort. Subjects that were awake for the entire nighttime scan were excluded (1 older from Site 2). Additionally, runs with average framewise displacement > 0.3mm or maximal framewise displacement > 2mm were excluded, resulting in the total exclusion of 3 young and 2 older subjects due to high motion in all runs. After exclusion, 25 young adults (age 20-39 years, mean = 24 years, 14 Female) and 39 older adults (age 61-84 years, mean = 68 years, 20 Female) remained in the analysis. 17 young and 21 older of the subjects analyzed were from Site 1 while 8 young and 18 older were from Site 2.

For the daytime experiments, 103 subjects in total completed an MRI session (36 young Site 1, 0 Young Site 2, 37 older Site 1, 30 older Site 2). There was overlap between subjects who completed daytime and nighttime experiments: 32 older and 20 younger subjects completed both experiments. For daytime experiments, subjects were allowed to consume their typical amount of daily caffeine that reported in their screening form the day of the scan. Because the daytime task stimuli were visual, subjects were screened for vision loss to minimize differences in vision across the two groups. Individuals in the young and older group were included only if they had contact- corrected non-pathological vision, or a prescription strength less than [–2.5, -2.5] diopters. A subset of daytime subjects were excluded due to equipment failure, visible motion artifacts in scans, or inability to stay awake during daytime task runs. After exclusion, the final daytime dataset consisted of 31 young (age 18-39 years, mean = 24.7 years, 16 Female) and 55 older subjects (age 60-85, mean = 69.7 years, 34 Female). In the final dataset, 59 subjects (31 young, 28 older) were scanned at Site 1, and 27 subjects were scanned at Site 2.

### Data Acquisition

#### MRI acquisition

Site 1: Subjects were scanned on a 3T Siemens Prisma scanner with a 64-channel head and neck coil. Stimuli were presented on a VPixx Technologies PROPixx Lite Projector (VPixx Technologies, Quebec, Canada) with a refresh rate of 120 Hz. Button presses to the visual task were recorded with a Current Designs 1x2 Button box. Eye movements were monitored with an EyeLink 1000 Plus Eyetracker (SR Research). Pulse oximetry, skin conductance, and respiration were monitored with BIOPAC sensors (BIOPAC Systems).

Site 2: Subjects were scanned on a 3T Siemens Prisma scanner with a 64-channel head and neck coil. Stimuli at site 2 were presented on a NEC NP4100 screen (size = 1024x768) at a 60 Hz refresh rate. Button presses were recorded with a Current Designs 932 Interface with the HHSC 2x2 button box. Eye movements were monitored with the Eyelink 100 Eyetracker (SR Research) with a long-range mount configuration. Pulse oximetry, skin conductance, and respiration were monitored with BIOPAC sensors (BIOPAC Systems).

Anatomical images were acquired using a 1 mm isotropic T1-weighted multi-echo MPRAGE (van der Kouwe et al., 2008). Functional runs were acquired using a gradient-echo EPI sequence (TR=0.378 s, TE=31 ms, 2.5 mm isotropic voxels, 40 slices, Multiband factor=8, blipped CAIPI shift=FOV/4, flip angle=37°, no in-plane acceleration).

#### EEG acquisition

At both sites, EEG was acquired using MR-compatible 32-channel EEG caps fitted with 4 carbon wire loops (BrainProducts GmbH, Germany) at a sampling rate of 5000 Hz. EEG acquisition was synchronized to the scanner 10 MHz clock to reduce aliasing of high-frequency gradient artifacts. Additional sensors were used to record systemic physiology: respiration was measured simultaneously using an MRI-safe pneumatic respiration transducer belt around the abdomen and pulse was measured with a photoplethysmogram (PPG) transducer (BIOPAC Systems, Inc., Goleta, CA, USA). Physiological signals were acquired at 2000 Hz using Acqknowledge software and were aligned with MRI data using triggers sent by the MRI scanner. Two built-in electrooculography (EOG) electrodes were used to record eye movements. Three electromyography (EMG) electrodes were recorded and one was selected for sleep stage scoring.

#### Daytime task design: Neurovascular coupling experiment

We presented a counterphase flickering checkerboard visual stimuli using Matlab and Psychtoolbox. Subjects viewed the stimuli with a mirror that was mounted on the 64-channel head coil. Stimuli flickered at a rate of 12 Hz. Each run lasted 360s, alternating between a fixed 16 s ON period and a variable OFF period (ranging from 30 s to 32.5 s). A red fixation dot at the center of the screen switched between dark and light red and subjects were asked to keep their eyes on this dot throughout the task and to press a button each time they observed a color change. Button presses and eye tracking videos were monitored during scans to confirm that subjects maintained wakefulness throughout the task. Two contrast conditions were collected for all subjects to control for possible differing contrast sensitivity across groups: a high contrast (90%) condition and a (45%) low contrast condition. All subjects were asked to complete a minimum of four runs of the visual task. A subset of subjects completed additional runs if time permitted.

#### Daytime task design: Cerebrovascular reactivity experiment

We presented instructions for the breathhold task using Matlab and Psychtoolbox. Subjects viewed the stimuli with a mirror that was mounted on the 64-channel head coil. The breathhold task consisted of eight one-minute trials. Each trial consisted of 1) three cycles of paced breathing with a 3-second inhale and 3-second exhale, during which the words “Breathe in” and “Breathe out” were displayed on the screen for the duration of the inhale and exhale periods, 2) a 3-second period during which “Prepare to hold your breath” was displayed on the screen, 3) a 15-second breathhold, during which a countdown timer on the screen displayed the remaining time left in the breathhold 4) one 3-second of exhale through the nose immediately after the breathhold, and 5) 21 seconds of recovery free breathing (‘Breathe freely’). Expired end-tidal CO_2_ gas was continuously measured through a nasal tube during the breathhold task using a CO_2_ analyzer (CapStar-100, CWE Inc., USA) that was pre-calibrated using a 5% CO_2_ gas tank. Signals were acquired using a PowerLab (PowerLab, AD Instruments) acquisition system. Subjects completed one run of the breathhold task.

#### Cerebrovascular reactivity (CVR) mapping analysis

CVR parameter maps were computed using a publicly available software package, *phys2cvr* version 0.18.6 (Moia et al., 2022), which implements a lagged-GLM framework that has been described previously (Moia et al., 2020). For preparing input data, we first spatially smoothed the slice-time and motion-corrected breathhold fMRI using a 2.5477 gaussian kernel (fslmaths, FSL version 5.0.9). Skull-stripped brain masks were computed using SynthStrip (Freesurfer version 7.4.1). Peaks were extracted from the end-tidal CO_2_ recording using *peakdet* version 0.3.0 (DuPre et al., 2024). The smoothed fMRI data, masks, and detected peaks were passed into phys2cvr which outputted CVR and lag maps. Any runs with a mean framewise displacement value above 0.3 mm were excluded, leading to exclusion of 2 subjects. We then excluded subjects based on previously published exclusion criteria (Zvolanek et al., 2023): the percentage of total power that is within a frequency band around the breathhold frequency [0.012, 0.022 Hz], taken from the end-tidal CO_2_ regressor signal (an output from peakdet software), was computed, and if that percentage is below 40% the data is excluded, leading to exclusion of 24 subjects. After applying exclusion criteria, 50 subjects remained. Voxels with negative CVR or with lag values above the boundary of the phys2cvr lag optimization (8.7 seconds) were excluded.

#### CSF ROI masking

CSF flow signals were measured from ROIs in the fourth ventricle. ROI masks were hand drawn on the single-band reference scans for each individual run, to account for potential motion between scans. The single-band reference image was used for ROI definition without any information from the functional runs, to remove any potential bias from selecting inflow voxels based on time-series information. To standardize the masks across subjects, the fourth ventricle was selected from the bottom four slices of the acquisition volume in all runs.

Nighttime analysis used the average of all voxels within this mask while the daytime analysis used only the first slice, which is most sensitive to inflow for analysis, because of the known smaller effect size of sensory-evoked compared to sleep-related changes in CSF flow.

### Data Preprocessing

#### EEG preprocessing

Gradient artifacts were removed using average artifact subtraction (Allen et al., 2000) with a moving average of the previous 20 TRs. Ballistocardiogram artifacts were removed from each EEG channel using signals from the 4 carbon wire loops using the sliding Hanning window regression method from the EEGLab CWL toolbox (van der Meer et al., 2016) with the following parameters: window=25 s and a lag=0.09 ms. Cleaned EEG signals were re-referenced to the average of EEG channels.

#### EEG sleep scoring

EEG data were processed according to AASM guidelines. EEG and EOG channels were filtered between 0.3-20 Hz. Two trained scorers manually sleep scored EEG data in 30-second epochs according to the AASM sleep scoring guidelines (Berry et al., 2017) using the *Visbrain* GUI (https://pypi.org/project/visbrain/). Any discrepancies between the two scores were discussed and merged before final analysis.

#### fMRI Preprocessing

All functional runs were slice-time corrected using FSL version 6 (slicetimer; https://fsl.fmrib.ox.ac.uk/fsl/fslwiki), (Woolrich et al., 2009) and motion corrected to the middle frame using AFNI (3dvolreg; https://afni.nimh.nih.gov/).). Physiological noise was removed using HRAN, open-source software that implements a statistical model of harmonic regression with autoregressive noise (Agrawal et al., 2020); https://github.com/LewisNeuro/HRAN). Motion corrected data were registered to each individual’s anatomical data using boundary-based registration (bbregister, (Greve & Fischl, 2009)).

#### Visual task EEG envelope calculation

Trial-averaged waveforms were calculated by averaging four posterior-occipital EEG channels (PZ, O1, O2, Oz) for each subject. Any trials with amplitude spikes greater than +50 uV or -50 uV were considered artifacts and were excluded from further analysis. Each subject’s trial averaged waveform was then bandpass filtered using zero-phase bandpass filtering between 11.99 and 12.01 Hz at the stimulus frequency (12 Hz). The amplitude envelope of the signal was extracted by taking the amplitude of the Hilbert transform. Statistical differences between the envelope amplitude of older and younger adults was tested with a Wilcoxon rank sum test across subjects within 8-second time bins, corrected with the Bonferroni-Holm method for multiple comparisons.

### Data Analysis

#### Daytime experiment: Visually-evoked BOLD response in primary visual cortex and gray matter

Masks for primary visual cortex (V1) and global gray matter mask were derived from the FreeSurfer recon-all parcellation (Fischl et al., 2004). All voxels within the mask were averaged within each region to produce a mean V1 BOLD signal, and a mean global gray (subcortical and cortical) matter signal. Signals were converted to percent signal change by dividing by the mean BOLD signal and were detrended to avoid low frequency artifacts related to scanner drift. The first 10 volumes were discarded to allow the signal to reach steady state. Trial-averaged BOLD signals were calculated for each individual in a window [-5 46] around the onset of the stimulus. Any trials with a framewise displacement value that exceeded 0.4 mm within the 50 second trial window were excluded from further analysis. All BOLD waveforms were low-pass filtered below 0.1 Hz. Statistical differences between the V1 BOLD responses to the visual stimulus across older and younger adults were tested with a Wilcoxon rank sum test within 8-second non-overlapping time bins, corrected with the Bonferroni-Holm method for multiple comparisons. Global gray matter stimulus-evoked BOLD magnitudes were calculated as the difference in mean BOLD signal during a peak window (t=3 to t=24), and a trough window (t=25 to t=35). These windows were chosen based on the timing of gray matter peak and trough waveforms reported in prior work (Williams et al. 2023). Group differences between stimulus-evoked global gray matter BOLD magnitudes were tested with an unpaired t-test across subjects.

#### Daytime experiment: fMRI analysis of two visual stimuli contrast conditions

fMRI BOLD and CSF flow data from runs with high contrast (90%) stimuli and lower contrast (45%) stimuli were averaged separately and compared with a Wilcoxon rank sum test corrected via Bonferroni Holm for multiple comparisons in a moving 8-second window. fMRI data were excluded for motion exceeding 0.4 mm, leading to exclusion of 4 older adults in the high contrast condition, and 5 older adults in the low contrast condition. All younger adults had both usable high and low contrast data available.

#### Daytime experiment: EEG analysis of two visual stimuli contrast conditions

The average envelope of EEG data from runs with high contrast (90%) stimuli and lower contrast (45%) stimuli were averaged separately and compared with a Wilcoxon rank sum test corrected via Bonferroni Holm for multiple comparisons in a moving 8-second window. Trials were excluded for amplitude exceeding 50 µV, leading to exclusion of 3 older adults in the high contrast condition, 1 older adult in the low contrast condition (total older low contrast n=52, high contrast n=54), and 4 younger adults in the high contrast condition, 5 young adults in the low contrast condition (total young adults: low contrast n=26, high contrast=27)

#### Daytime experiment: Awake visually-evoked CSF inflow signal

For analysis of daytime CSF data, CSF inflow signals were extracted using only the first slice of the four-slice hand-drawn fourth ventricle masks. In contrast to sleep, during wakefulness, CSF flow velocities are much lower (Williams et al., 2023) and therefore typically produce flow signals limited to the bottom-most or bottom two slices due to the known effect of flow velocity on fMRI signals (Gao et al., 1996). Therefore, for our daytime data, we used only the bottom slice of the mask for wakefulness inflow signals. To standardize the number of voxels used across subjects, we selected the two voxels from the bottom slice for each run that contained the largest CSF inflow signals.

To create an individual-level metric of stimulus-evoked CSF magnitude, we calculated the difference in CSF magnitude during expected trough and peak windows. As CSF flow alternates with blood fluid, we hypothesized CSF to be suppressed after stimulus onset during the ON block, when the BOLD response peaks. Conversely, we hypothesized maximal CSF flow during the OFF block, when the BOLD decreases to baseline, allowing CSF to flow upwards into the brain.

#### Nighttime experiment: Spectral power analysis

The power of fMRI and EEG signals was calculated using multitaper spectral estimation in Python using the spectral_connectivity toolbox (Denovellis et al., 2022). Power was estimated in 60-second segments of continuous wakefulness or NREM sleep, and any 60-second segments that had average framewise displacement > 0.3 mm or maximal framewise displacement above 0.75 mm were excluded from analyses. For the fMRI and CSF analyses 5 tapers were used. For EEG multitaper spectral estimations 25 tapers were used. We computed the power spectrum in each electrode independently and excluded channels that were identified as bad at the EEG preprocessing stage by visual inspection of the spectrogram on a run-by-run basis. Additionally, segments from any channels that had a maximum amplitude over 150 µV were excluded. The BOLD, CSF, and EEG spectra were averaged across individuals within each group during wake and NREM sleep separately, and the standard error was calculated across subjects. Both the low-frequency (0.01–0.1 Hz) CSF and BOLD power as well as the frontal EEG delta (0.5–4 Hz) power were calculated as the sum of power in the respective band and then converted to decibels. Analyses comparing frontal EEG delta across groups used the Fz electrode; in runs with bad quality signals ini Fz, we instead used F3 (n_subjects_=7, n_runs_=10). Group-level comparisons between CSF, BOLD, and frontal EEG delta power were performed using the Wilcoxon rank-sum test across subjects. Pairwise comparisons across sleep and wakefulness were performed in subjects that had both wake and NREM sleep data (n = 32 older, 21 younger) using a paired t-test. Group-level differences in the BOLD spectra were calculated using the Wilcoxon rank-sum test and were Holm-Bonferroni corrected for multiple comparisons.

#### Nighttime experiment: Cross-correlation analysis

We extracted 90-second segments of continuous NREM sleep from all participants and excluded any segments with average framewise displacement above 0.3mm or maximal framewise displacement above 0.75mm. BOLD and CSF signals were filtered between 0.01–0.1 Hz. The EEG signal from Fz (or F3) was filtered between 0.2–4 Hz using a finite impulse response filter. The delta amplitude was calculated as the magnitude of the Hilbert transform and was then smoothed using a 4-second moving average. The cross-correlation within each segment was calculated and then averaged within groups. A permutation bootstrap test with 10,000 iterations was performed to compare the strength of the correlation between the two groups.

#### Nighttime experiment: CSF peak locked analysis

We identified peaks in the CSF signal during sleep by first converting the signal to percent signal change using the 15th percentile as the baseline, then filtering the CSF signal between 0.01–0.1 Hz, and identifying local maxima that surpassed a threshold of 5% amplitude. These peaks were then used to extract a peak-locked signal of the frontal EEG delta (0.2–4 Hz) amplitude envelope. To test for significant differences in the peak of the delta envelope we performed a permutation test on the difference in delta peak amplitude between young and older adults using 10,000 iterations.

#### Nighttime experiment: Mediation analysis

Mediation analysis was conducted in R using the *lavaan* package (Rosseel, 2012). The choice of a serial mediation model was based on a prior model of the link between EEG and CSF (Fultz et al., 2019). The predictor variable (X) was coded a categorical variable. The other variables, frontal delta power (M_1_), low-frequency gray matter BOLD power (M_2_), and low-frequency CSF power (Y), were obtained as described above in the *Spectral Power Analysis* section. Confidence intervals for coefficients were obtained by bootstrapping and the p-values were obtained under the Chi-squared distribution.

## Acknowledgments

We are grateful to Courtney Zambello for project management support, and to Emilia Schimmelpfennig, Zenia Valdiviezo, Shruthi Chakrapani, Dr. Stephanie McMains, and Steve Shannon for assistance with experiments. This work was funded by National Institutes of Health grants R01-AG070135, R01-AT011429, U19-NS128613, the Simons Collaboration on Plasticity in the Aging Brain (811321), the McKnight Scholar Award, the Pew Biomedical Scholar Award, and the Sloan Fellowship.

---

**Supplementary Figure 1:**
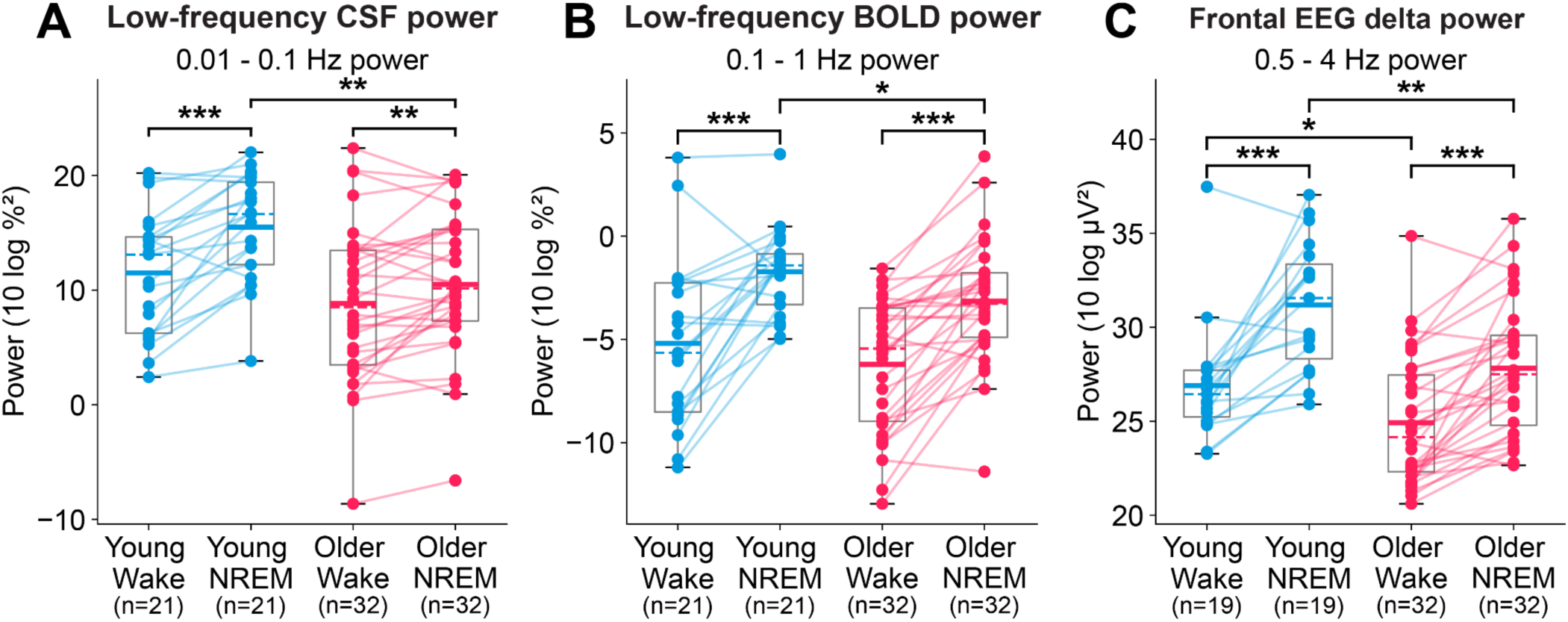
Paired analysis of CSF, BOLD, and EEG bandpower in subjects with both sustained wakefulness and sleep. **A)** Low-frequency (0.01-0.1 Hz) CSF power significantly increases from wakefulness to NREM sleep in both groups (p_young_<0.001, p_older_=0.007, paired t-test). Additionally, low-frequency CSF power during NREM sleep is significantly lower in older adults compared to young adults (p=0.001, Wilcoxon rank-sum test). **B)** Low-frequency gray matter BOLD power significantly increases from wakefulness to NREM in both groups (p_young_<0.001, p_older_<0.001, paired t-test). Similar to CSF, low-frequency BOLD power during NREM sleep is significantly lower in older adults compared to young adults (p=0.03, Wilcoxon rank-sum test). **C)** Frontal EEG delta power significantly increases from wakefulness to sleep in both groups (p_young_<0.001, p_older_<0.001, paired t-test). Frontal delta power is significantly reduced in older adults during both wakefulness and sleep (p_wake_=0.025, p_sleep_=0.003).

**Supplementary Figure 2:**
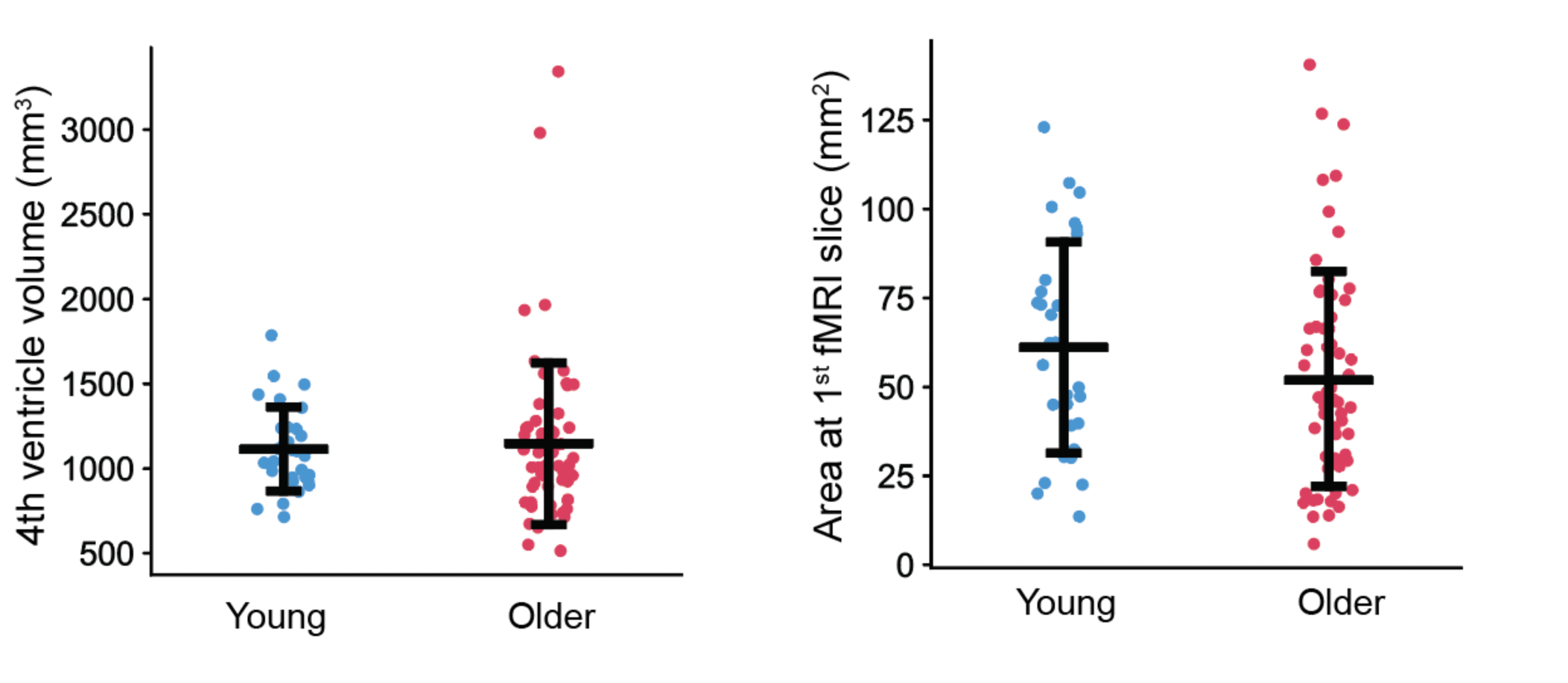
Fourth ventricle area and volume does not significantly differ across age groups. **A)** Older adults (pink) did not have significantly different fourth ventricle volumes than younger adults (blue). **B)** Older adults also did not have significantly different cross-sectional areas at the edge of the fMRI volume intersecting the fourth ventricle, where CSF inflow signals were measured. These control analyses confirm age-related CSF inflow differences were not due to age-related changes in fourth ventricle size or positioning.

**Supplementary Figure 3.**
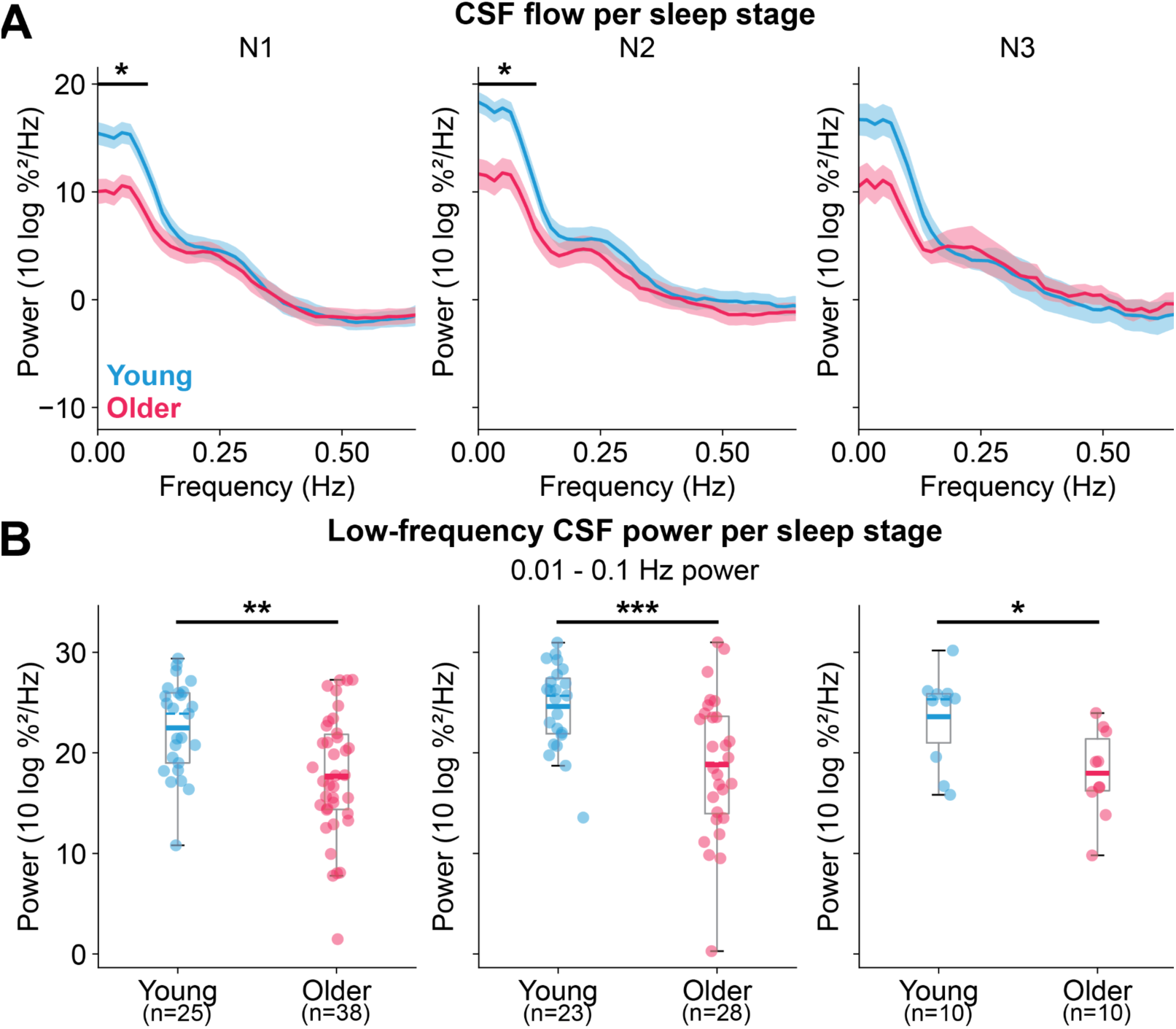
CSF spectra and low-frequency power per NREM sleep stage. **A)** CSF power spectra during N1, N2, and N3 NREM sleep in both groups. During N1 and N2 young adults had significantly higher power in the [0, 1.11] band (p<0.05, Wilcoxon rank-sum test, Holm-Bonferroni corrected). **D)** Total [0.01, 0.1] Hz CSF flow power is significantly higher in young adults compared to older adults in all 3 NREM stages (p_N1_=0.002, p_N2_=0.001, p_N3_=0.01, Wilcoxon rank-sum test).

**Supplementary Figure 4.**
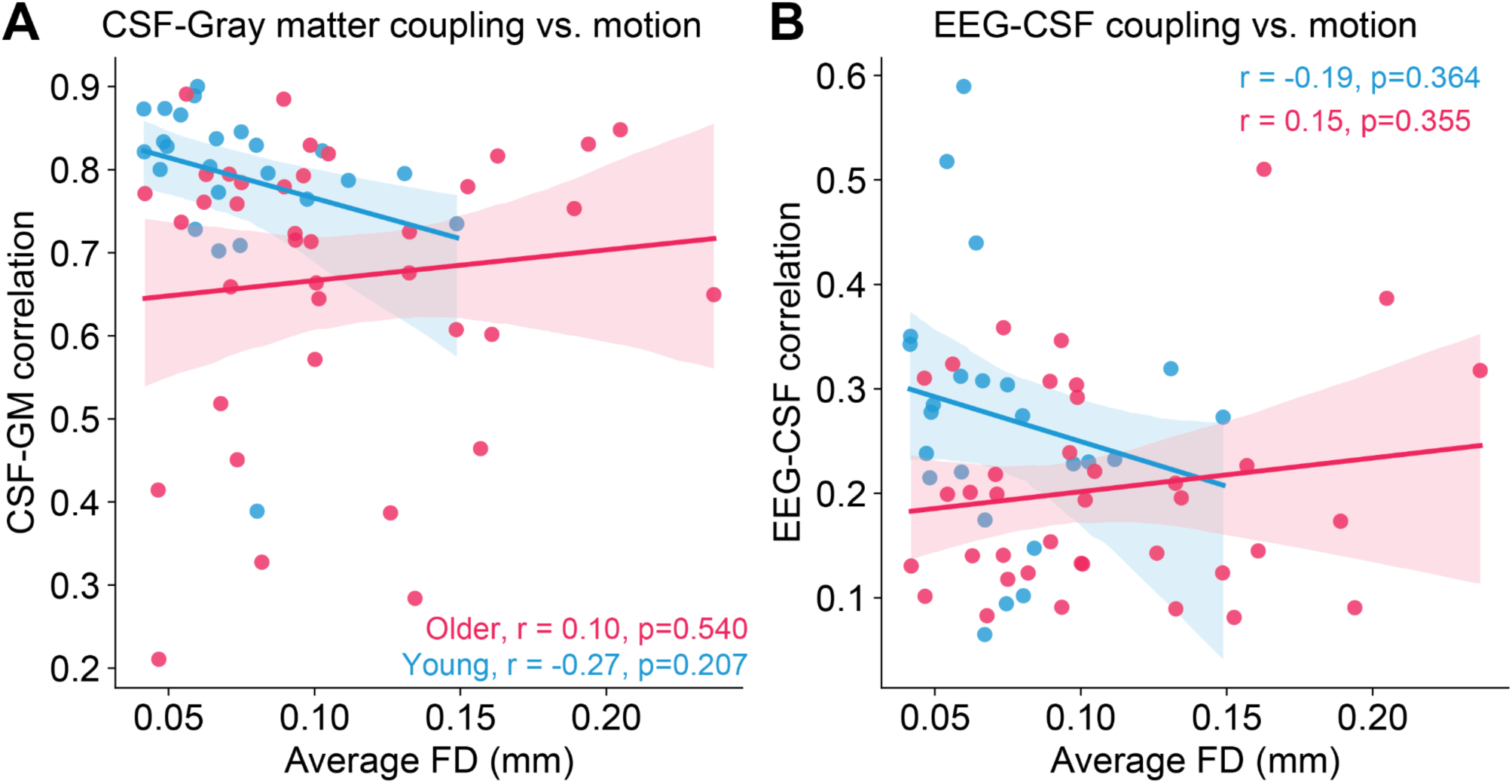
Framewise displacement is not significantly correlated with the strength of coupling between CSF and gray matter or between CSF and EEG. **A)** Average framewise displacement during segments used to calculate correlation between CSF and gray matter is not significantly predictive of the strength of correlation in either group (r_young_=-0.27, p_young_=0.207, r_older_=0.10, p_older_=0.540). **B)** Average framewise displacement during segments used to calculate EEG and CSF correlation is not significantly related to the strength of correlation in either group (r_young_=-0.19, p_young_=0.364, r_older_=0.15, p_older_=0.355).

**Supplementary Figure 5.**
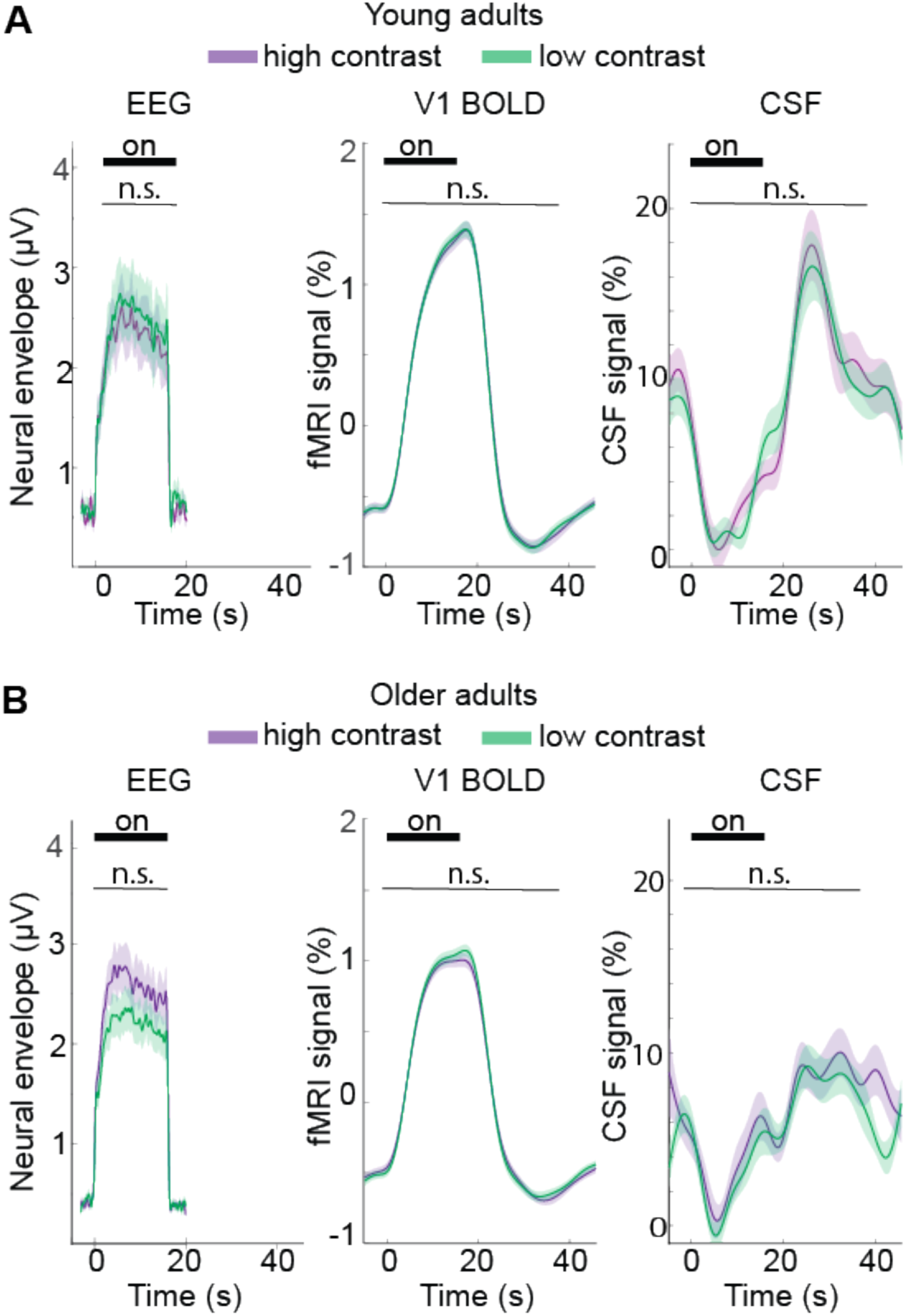
Visual contrast sensitivity control experiments show no difference in response to high and low contrast conditions of daytime visual stimuli. **A-Left)**- Waveforms are mean EEG envelope amplitudes in young adults to the high contrast (90%, n=27, purple) condition and the lower contrast (45%, n=26, green) condition. A black bar indicates the duration of the ON visual stimulus block. **A-Middle**) Waveforms are the mean v1 BOLD responses in the young group to daytime visual stimuli. Colors indicate high (n=31, purple) and low (n=31, green) contrast conditions. **A-Right)** Waveforms are group mean pulses of evoked CSF inflow in the young group to the two contrast conditions. **B-Left)** Waveforms are mean EEG envelope amplitudes in older adults to the high contrast (n=52, purple) condition and the lower contrast (n=54, green) condition. **B-Middle)** Waveforms are the mean v1 BOLD responses in the older group to high (n=51, purple) and low (n=50, green) conditions. **B- Right)** Waveforms are the group mean pulses of evoked CSF flow in the older group to high (n=51, purple) and low (n=50, green) contrast conditions.

**Supplementary Figure 6.**
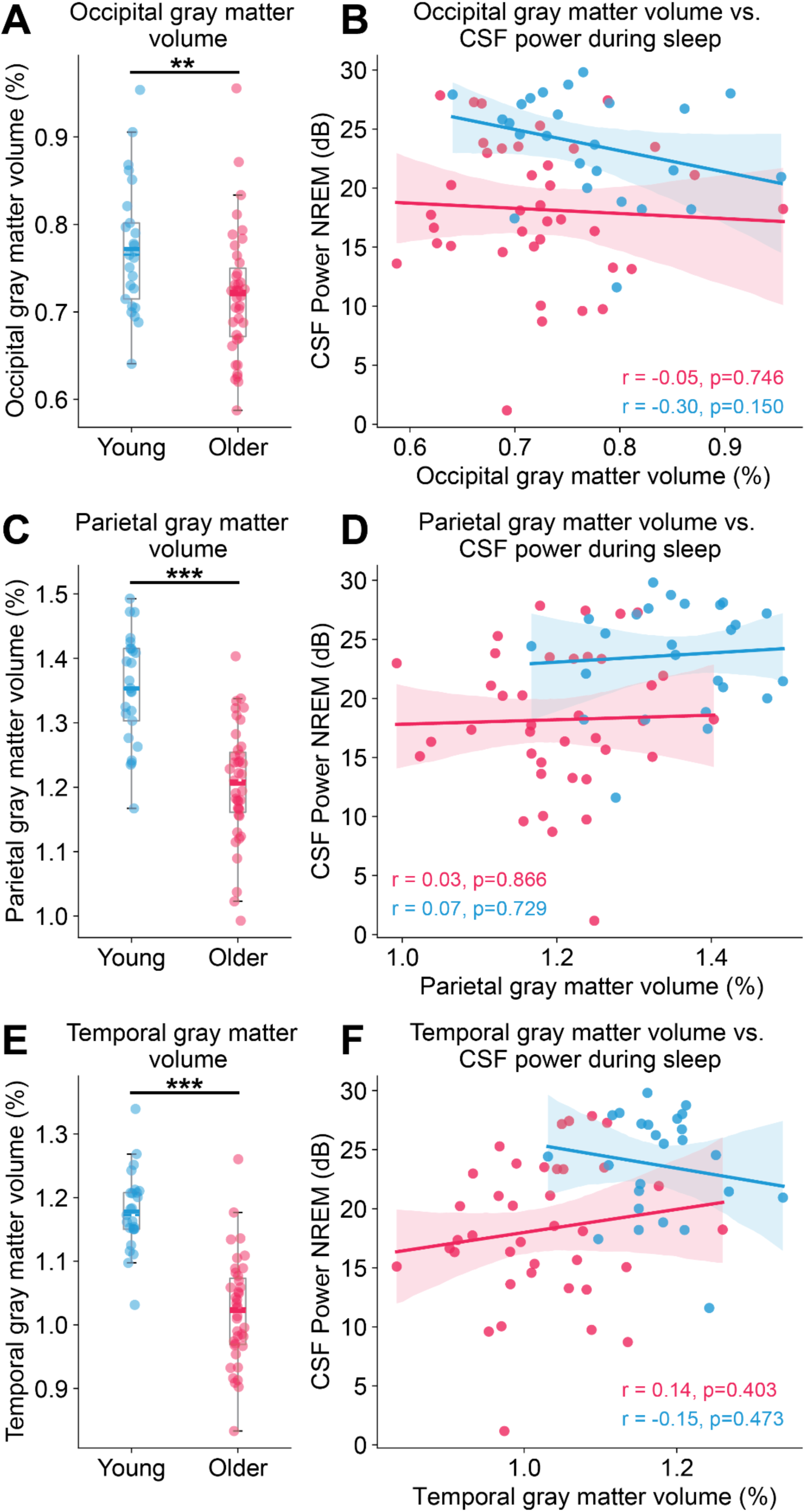
Occipital and parietal lobe gray matter do not exhibit the same significant relationships with CSF flow as frontal gray matter. **A)** Older adults had significantly less gray matter in the parietal lobe normalized to total intracranial volume (p<0.001, Wilcoxon rank-sum test). **B)** Low-frequency CSF power during NREM sleep is not significantly associated with normalized parietal lobe gray matter volume in either age group (r_older_=0.03, p_older_=0.866, r_young_=-0.07, p_young_=0.729). **C)** The strength of coupling between CSF and EEG is not significantly related to normalized parietal lobe gray matter volume (r_older_=0.13, p_older_=0.445, r_young_=0.11, p_young_=0.612). **D)** Older adults had significantly less gray matter in the occipital lobe normalized to total intracranial volume (p = 0.007, Wilcoxon rank-sum test). **E)** Low-frequency CSF power during NREM sleep is not significantly associated with occipital lobe gray matter volume normalized to intracranial volume in either age group (r_older_=-0.05, p_older_=0.746, r_young_=-0.30, p_young_=0.150). **F)** The strength of coupling between CSF and EEG is not significantly related to normalized occipital lobe gray matter volume (r_older_=0.01, p_older_=0.970, r_young_=-0.05, p_young_=0.803).

## Notes

### Competing Interest Statement

The authors have declared no competing interest.

